# Diffusion models learn underlying trends in actomyosin networks and predict behavior at unseen filament turnover

**DOI:** 10.64898/2026.05.26.727950

**Authors:** Elisabeth Rennert, Agnish Kumar Behera, Yuqing Qiu, Suriyanarayanan Vaikuntanathan

## Abstract

Generative diffusion models have demonstrated an ability to produce novel images sampled from the learned underlying data distribution. These models are able to infer system characteristics for parameter combinations that were not seen during training. We investigate the ability of these models to infer trends in biological data from limited samples. Specifically, we consider the response of system scale behaviors such as cortical flow in a simulated actomyosin system as we tune filament turnover rates. We train a diffusion model on coarse grained actin curvature and density heatmap images, and are able to generate images from conditioning variables not seen during training. These images are predictive of nonlinear trends in the system. We also consider characteristics of the system that allows this level of inference, such as the strong linear relationship between average density and filament turnover in the system, and explore minimal underlying dynamics with a motor binding model.

## I. INTRODUCTION

The actin cortex is crucial to the generation and transmission of forces within a cell. This actively assembled biopolymer network interacts with a variety of proteins that modulate the network structure and growth [1]. The balance between active stresses generated by myosin motors and network viscosity can produce coordinated and sustained flows in the actin cortex [2, 3]. Cortical flows establish polarity within the cell to direct motility [4, 5], cytokinesis [6] or morphogenesis [7–9]. This wide range of applications is possible due to the highly tunable properties of actin networks, partly controlled by their regulatory interactions with various polarity establishing proteins.

Cortical flows are implicated in the establishment of polarity in *C. elegans* embryos through asymmetrical contractions establishing a localization of PAR proteins to the developing anterior pole of the cell [7]. The patterning protein CDC-42 creates a gradient of the molecular motor myosin which drives the cortical flow toward the anterior half of the cell [10, 11]. The ability to predict cortical flows and their impacts on cellular processes across parameter perturbations is currently limited by the availability of detailed observations of the actin cortex’s structure and dynamics, whether through experiments or detailed, and thus computationally expensive, simulations. Given the high level of network variability that can be produced even from a limited set of components, an ability to predict mesoscale system behaviors from limited samples would allow more rapid exploration of high dimensional parameter spaces. Generative diffusion models have emerged in recent years as powerful tools to create new samples from complex high dimensional distributions [12–17]. In this work we investigate the ability of these models to generalize across the parameter space and predict cortical flow in novel regimes.

Experiments in reconstituted actin networks [18] and in *C. elegans* embryos [11] have shown that the plastin, formin, and profilin concentrations can non-monotonically tune the rate of cortical flow, suggesting a mechanism to control flows in embryo development. Across perturbations, the filaments show a consistent rate of nucleation with distinct assembly and disassembly phases. We will describe this actin assembly/disassembly rate as the turnover throughout this paper. This system’s cortical flow behavior in response to varying turnover can be reproduced through carefully tuned Cytosim simulations [3, 11, 19]. In these simulations, there is a non-monotonic response of cortical flow with increasing turnover, shown in Fig. 1c. Our main result shows how generative diffusion models are able to predict non-monotonic trends just from static images provided at points from different mechanical regimes, suggesting a capacity to identify underlying interactions and predict resulting system scale dynamical behaviors like cortical flow.

**FIG. 1.**
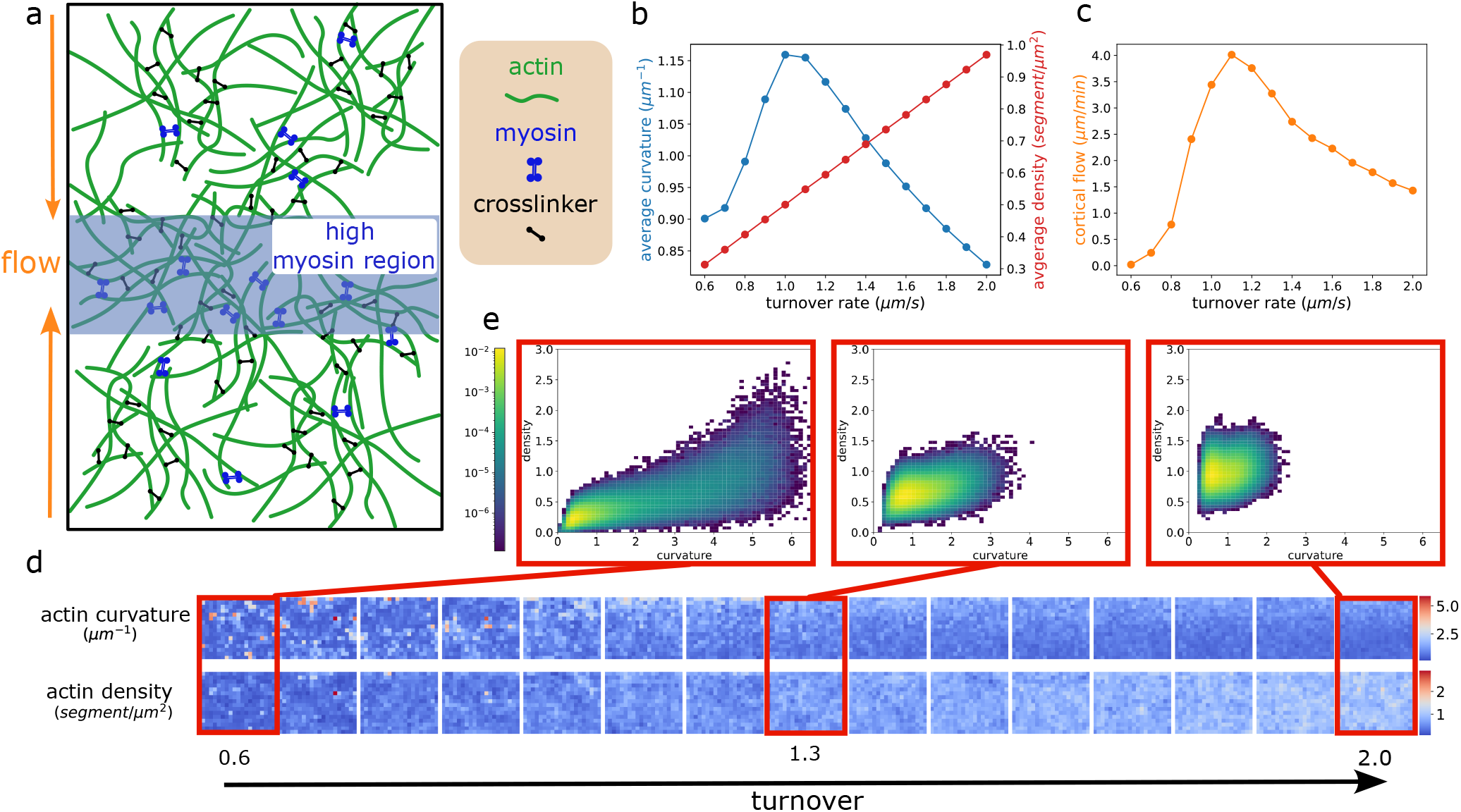
(a) A schematic of an actomyosin system containing actin, myosin and crosslinkers. The high myosin region in the center creates a myosin gradient which drives cortical flow towards the center of the simulation box. See SI for simulation details. (b) Trends in average actin curvature and density show a linear density-turnover relationship and a peak in the curvature. Density is calculated as the number density of 0.2 *µ*m filament segments in the simulation box. (c) The average rate of cortical flow of actin towards the center of the box is dependent on the ratio of active stress and viscosity, which itself depends in part on the local curvature and density. (d) Heatmap data used for training. Curvature and density fields are calculated by binning the simulation data and mirroring over the center line (so the top edge is the highest myosin concentration). (e) Joint distributions of curvature and density pixel intensity in log scale. Distributions are across all pixel locations, removing spatial information.

## II. METHODS

### A. Coarse grained simulation data captures system scale trends

The specific simulation setup we consider is a 2D system of filamentous actin, myosin and crosslinkers with a high myosin region at the center, as shown in Fig. 1a. The imposed myosin gradient causes anisotropic contractility which drives flows of actin [9] towards the center of the box. We employ the turnover dynamics described in [3], where the filament nucleation rate is fixed and the filaments grow for a fixed time at the specified turnover rate before disassembling at that same turnover rate. We simulate this system using the Cytosim [19] simulation package at 15 different actin turnover rates. This configuration generates consistent cortical flows towards the center of the simulation box, in agreement with experimentally observed trends during polarity establishment in the *C. elegans* embryo [11].

To prepare the training data for our diffusion model, we reduced the full simulation data to coarse grained fields that are still predictive of network responses to perturbations. Analysis of autoencoder latent space suggests that both the density and curvature fields are required to predict flows [3]. Additionally, local actin curvature and density can also be measured experimentally through video microscopy, making the ability to infer trends from these fields broadly relevant. We create heatmap images of the actin curvature and density fields by dividing the simulation box into 1 *µ*m x 1 *µ*m bins, averaging the simulated network values within those bins, shown in Fig. 1d.

Due to the fixed actin growth window, the turnover rate determines the length of the filaments in the system. In combination with the constant nucleation rate, this results in the strong linear relationship between turnover and actin density shown in Fig. 1b. The peak in curvature is due to a decrease in network mesh size with increasing actin density. A smaller mesh size means the motors are more likely to bind a filament, exhausting the available pool of motors. After the motors are mostly bound, increasing actin density results in a drop in density of bound motors per length of assembled actin. Both the cortical flow and average curvature exhibit a peak as the turnover rate increases, with the curvature peak corresponding to the point with the highest motor density on the filaments. This is indicative of an underlying shift in the motor binding regime. This transition can also be seen in the shift in shape of the curvature-density joint distribution (Fig. 1e), with a much longer tailed curvature distribution at low turnover, shifting to a more compact distribution at high turnover. The long tail in the curvature distribution at low turnover indicates more extreme fluctuations in the relatively lower amount of F-actin.

If a generative diffusion model is able to infer the non-linear curvature trend in the system arising from an underlying shift in motor activity, that indicates an ability to generalize across mechanical paradigms that could allow it to predict more complex behaviors like cortical flow.

### B. Diffusion models conditioned on physically relevant parameters

We train a score-based generative model (SGM) to learn a score function in the space of actin heatmap images conditioned with the actin turnover rate. During training our model learns a conditional score function given by **S**(**x**_*t*_, *c*) = ∇ log *P*_*t*_(**x**_*t*_|*c*), where *c* is the class label, in our case the turnover rate. Our goal is to sample generated heatmaps conditioned with turnover rates not seen during training. We considered a variety of models trained on subsets of the 15 simulated turnover rates. For these models we adjust the number of frames seen per class in training such that all models are trained on 54,000 total images.

A physically relevant continuous conditioning variable allows us to avoid the problem of identifying an embedding that preserves the semantic relationships between class labels as is necessary with natural language descriptions, though it limits us to sampling heatmaps along that axis. As the turnover rate increases the resulting network behaviors, and thus underlying distributions of the heatmap images, shift. The high levels of overlap in distributions across turnover rates indicate that an individual turnover rate might contain information about neighboring rates. The conditional score functions learned by diffusion models capture the relationship between the data and conditional labels to the extent they can even be used as a classifier[20]. This indicates that the conditional score functions contain some core features of how the data varies across classes, and might pick up on underlying relationships and trends.

The structure and complexity of the training data has a strong impact on the types of models that are most effective and the level of prediction that can be drawn from the model. With this work we focus on the coarse-grained heatmap data rather than the full simulation as we are concerned with the ability of the model to infer nonlinearity in the large scale trends, rather than making specific predictions of microscopic structures. The heatmap data has only limited spatial anisotropy, as can be seen in Fig. 1d, with areas of high curvature slightly more likely at the top of the images due to the myosin gradient. This means the primary challenge is in capturing both the averages and the extreme fluctuations in the system. The coarse grained nature of the data combined with the high levels of overlap between different simulated turnover rates indicates there is not much hierarchy in the features the model learns, so the diffusion trajectories do not speciate (Fig. 2a) into separate distributions as is seen in more hierarchical data [21] like photographs. This allows us to use the simplest version of a SGM without noise scheduling or high levels of tuning and still learn characteristics of the underlying data distributions. We learn more general relationships within the data that indicate system level trends than details of the microscopic interaction of the individual components.

**FIG. 2.**
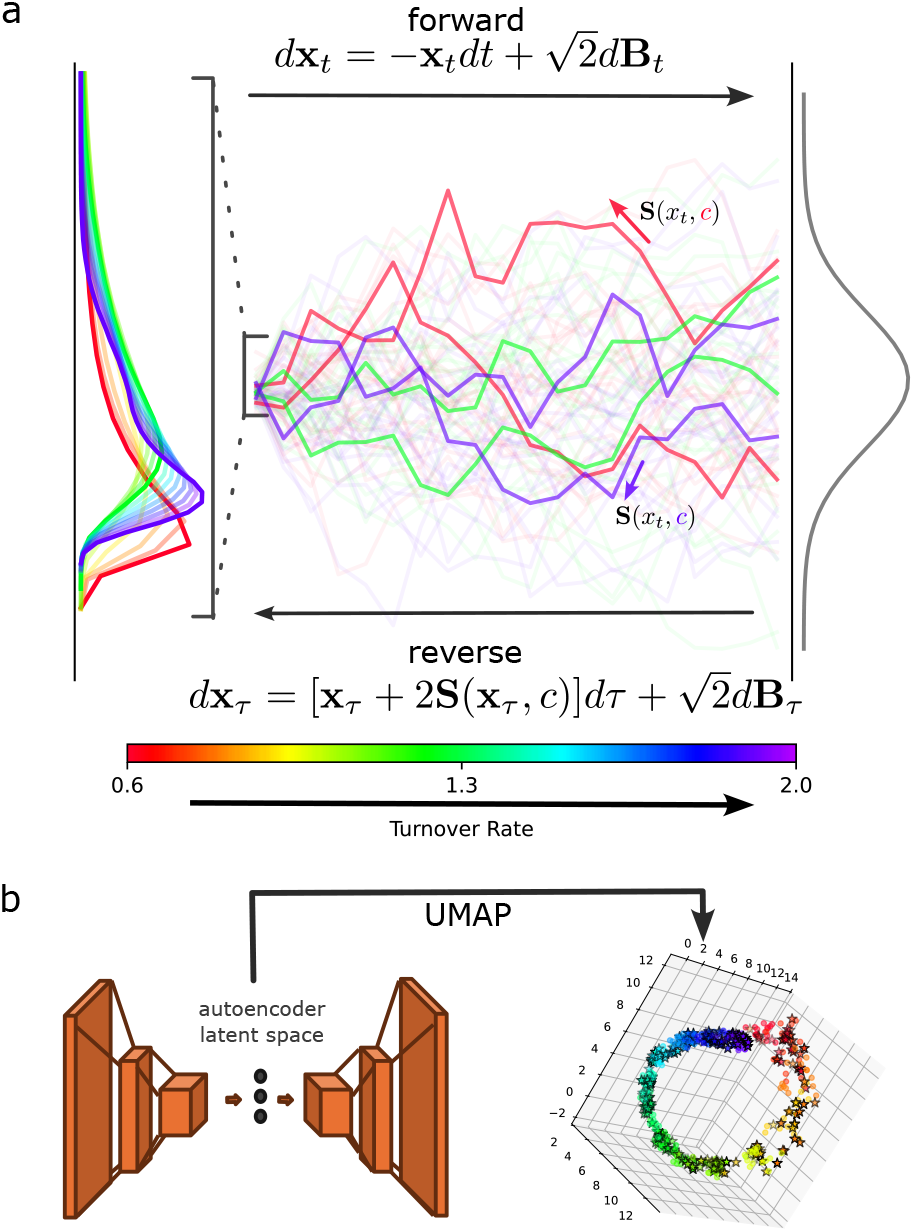
(a) Schematic of trajectories mapping between the turnover dependent data distribution and a Gaussian distributed prior. We train a SGM to learn a score function conditioned on turnover rate that can sample heatmap images from the specified rate. The turnover rate acts as a control parameter rather than a discrete class label, so the conditional data distributions have significant overlap. Distributions shown are of curvature pixel value across all spatial locations. (b) A convolutional autoencoder is trained on the simulated heatmap data. The autoencoder uses 3 convolutional downsampling layers to reduce the heatmap images to a 40 dimensional bottleneck. We use a UMAP of the low dimensional latent space to visualize the data and compare the overlap in latent distributions.

While we train on static heatmap images without the temporal correlations necessary to calculate cortical flow, there are other machine learning methods that attempt to predict the dynamic behaviors of such networks from static snapshots [22–24]. Other approaches to predict cortical flow could include a hydrodynamic model for this specific system configuration that connects the curvature and density fields to cortical flow. Additionally the model could be trained on sequences of heatmap images to also learn the temporal correlations that reflect the cortical flow in the system.

To evaluate the quality of the generated images, we train a convolutional autoencoder on the heatmap data. This model has a 40 dimensional bottleneck which provides a representation of key features necessary to reconstruct the data. We further reduce the dimensionality by taking a UMAP projection of the autoencoder latent space to visualize the heatmap distributions in 3D as shown in Fig. 2b. In both the simulated and generated heatmaps, the images are distributed along an underlying manifold corresponding to the turnover rate, with some overlap between neighboring rates, indicating this projection largely captures the high level features of the data. This further allows us to evaluate the overlap of the latent image distributions between simulated and generated heatmap images.

## III. RESULTS

Our model is able to successfully predict the global trends in average density and curvature, even across the nonlinearity in curvature as the system shifts motor binding regimes. For a model trained on the 6 extreme turnover rates (the 3 highest and 3 lowest turnover values) of our simulation, it was able to generate images that followed the average curvature and density trends from the full simulation as shown in Fig. 3a. The model could still predict the nonlinearity with as few as 4 seen turnover rates. While a naive spline interpolation was only slightly worse at predicting average trends with 6 seen rates, at 4 rates the generative model had substantially better predictions, shown in SI Fig. S2. Furthermore, a spline interpolation provides information only about average values without predicting any qualities of the distribution. The latent distributions in the autoen-coder bottleneck were also in agreement as can be seen in the UMAP visualizations in Fig. 3b, indicating enough identifying features were created in the generated images to target them to the correct areas in the autoencoder latent space. There can be some tradeoffs in the generated images between matching the averages and fully capturing the long tailed nature of the true distributions at lower turnover rates. The generated images tended to be more Gaussian in distribution, which is not surprising given the Gaussian prior of the diffusion model.

**FIG. 3.**
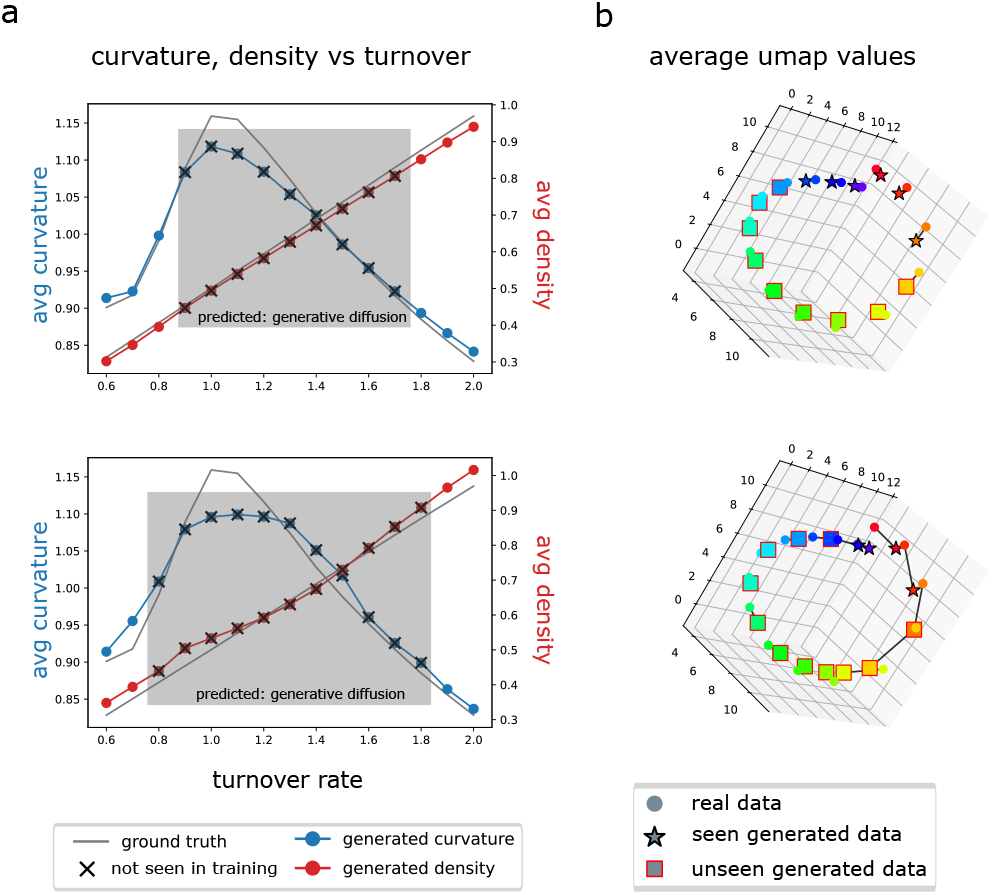
Diffusion models are able to infer nonlinear trends in system level characteristics across varying gaps in training data. (a) Comparison of generated vs simulated (ground truth) average curvature and density with increasing turnover rate. Results are shown for the 6 (top) and 4 (bottom) extreme turnover rates seen in training. Black x’s correspond to unseen turnover rates. The model is able to capture the curvature peak with only a few turnover rates at each extreme seen during training. (b) UMAP visualization of autoencoder latent space of heatmap data. Points are averages for the turnover rate indicated by the color, with shape indicating the simulated or generated data. Closer overlap indicates higher similarity between the simulated and generated images.

The size of the gap in turnover rate significantly affected the ability of the model to predict behaviors. Training on 4 turnover rates seems to be the minimum number of turnover rates and largest prediction gap size that can give accurate results. However, there can still be a lot of variation in model accuracy both during training and between models trained on the same subsets of data. This indicates there could be numerous spurious minima in the high dimensional loss landscape that the model can fall into without fully learning the details of the distribution. The model variability can be overcome by training an ensemble of models and choosing the ones with closest overlap between simulated and generated images at the training turnover rates.

The ability of these models to generate image data that follow underlying physical trends suggests a further ability to infer dynamic behaviors like directly predicting cortical flow with the right model architecture and resolution of data. The complexity of what the model needs to garner from the data is also demonstrated in that the limit in the model’s ability to infer behavior is not completely defined by the amount of data or even the turnover gap size, but also the number of distinct turnover rates seen in training.

## IV. DISCUSSION

Given the ability of the model to predict characteristics of the actomyosin system at unseen turnover rates, we consider what features of the data allow the model to make these predictions from limited data. The lack of tuning required for the model and the simplicity of the underlying SDE indicate the existence of some minimal underlying dynamics driving these trends.

The quality of the predictions we have shown indicates that there is information about other unseen turnover rates contained in the training data. We propose that the linear relationship between turnover and density allows the model to map fluctuations away from the mean at seen turnover rates to behaviors at unseen rates. This is aided by the high degree of overlap in the distributions, even across the full range of turnover rates considered. In Fig. 4b, we show slices from the joint distribution at the average densities associated with various turnover rates in the simulation data. The shape of these slices changes as the density approaches the point corresponding to the curvature peak, and the mode shift in motor saturation. These slices reflect how the trends across turnover rate are linked to density fluctuations within the network. Even at very low turnover rates, and thus low average densities, there are still some areas of the network experiencing a high enough actin density that a different motor binding regime can be observed. That might allow the diffusion model to learn the densities at which these shifts occur with relation to the conditioning variable and accurately reflect the curvature nonlinearity in the generated images. The slices also reflect the extent to which these fluctuations can be effective at predicting behavior, as the amount of information in the slices drops off with more extreme fluctuations.

**FIG. 4.**
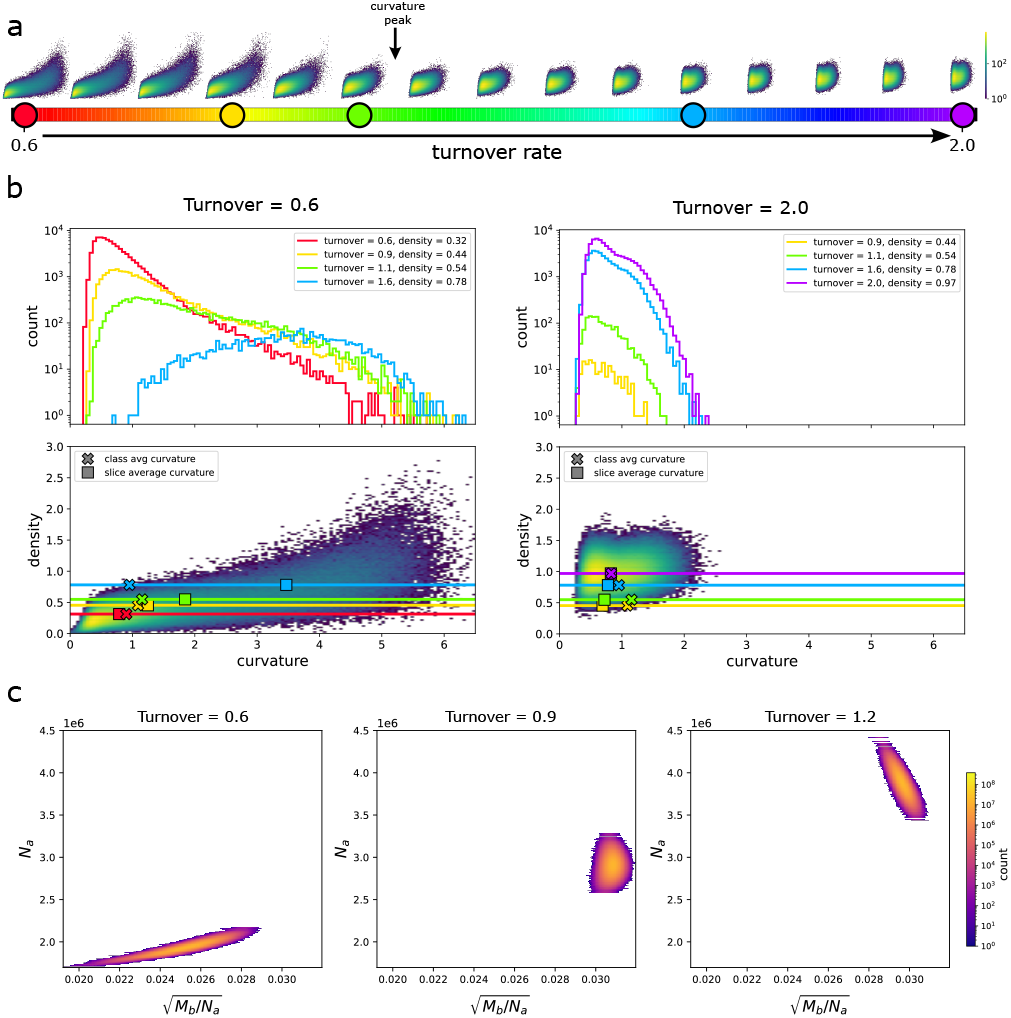
The joint distribution of curvature and density of the simulation data contains information about the distributions at higher turnover rates. (a) shows the joint curvature density distributions for all simulated turnover rates. The shape of the distribution shifts from a long tailed distribution at low turnover to a more compact distribution after the peak. Circles indicate the turnover rates corresponding to the slices taken in (b). (b) The joint curvature-density distributions at the 2 extreme turnover rates are considered. Slices are taken at the densities corresponding to the average density of the turnover rate indicated by the color. (c) Joint distributions produced by the minimal motor binding model. The model captures the shift in correlation as turnover increases and the trends in average values, though they have substantially less variance than simulated joint density and curvature distributions, leading to less overlap between individual turnover rates.

### A. Minimal model of motor binding regimes

If the diffusion model is picking up on system wide trends in the fluctuations to predict different regimes, then there is likely some characteristic underlying interaction giving rise to these trends. We consider a model for the number of bound motors, *M*_*b*_, from a pool of *M* total motors, with the rate equation given by

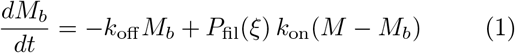

where *k*_on_ and *k*_off_ are motor binding and unbinding rates, and *P*_fil_ is the probability that the motor will diffuse into a filament during its lifetime.

This probability is given by

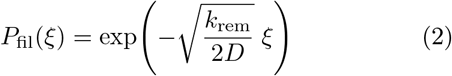

where *k*_rem_ is the motor removal rate that determines motor lifetime, and *D* is the motor diffusion constant. *P*_fil_ varies based on the mesh size *ξ* of the actin network. Network mesh size is deeply connected to the details of the crosslinking within the system, which we are not considering in this model. However, in quasi-2D actin networks, the mesh size has been described as scaling with actin concentration generically as [*c*_*A*_]^−1*/*2^ [25, 26]. We will use 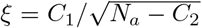, with *C*_1,2_ as fitting parameters to tune the mesh size scaling, where *C*_1_ sets the scale of the mesh sizes, corresponding to the mesh size at the minimum turnover considered, and *C*_2_ is an offset related to the minimal actin density needed to create a mesh. However these variables are approximate, so we fit them to get the best agreement between the normalized curves of 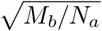 and the simulated average curvature. *N*_*a*_ is the number of assembled actin monomers in the system, proportional to the actin density. *N*_*a*_ fluctuates independently of the number of bound motors, distributed as a compound Poisson process in the number of filaments *N*_*f*_ and filament length *l*,

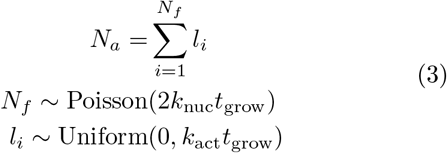

The distributions of *N*_*f*_ and *l* are a result of the specific actin growth dynamics in our system, depending on the filament nucleation rate *k*_nuc_, the filament growth time *t*_grow_, and the turnover rate *k*_act_. See SI for details.

We can then sample the joint fluctuations in *N*_*a*_ and *M*_*b*_ by sampling a value of *N*_*a*_ and using that to determine the Markov transition rates in motor binding, which we can use to sample *P* (*M*_*b*_|*N*_*a*_). This model describes the fluctuations in assembled actin and bound motors in the system, but to compare to the joint distributions of the heatmap data we need to connect actin curvature to bound motor density. If we assume the filament bending energy can be approximated by a worm-like chain, and each motor makes a unit energy contribution to the bending of a local segment, the curvature is approximately proportional to 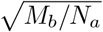.

While the joint distributions the model produces (Fig. 4c), do not fully capture the fluctuations of the simulation distributions, they do reflect a shift between 2 distinct motor binding regimes with increasing turnover. As the pool of available motors is exhausted, the fluctuations in the system become dominated by density fluctuations. The model may begin to diverge from the Cytosim simulation more at very high turnover rates as bundling effects, which were not considered in the model, become more impactful. This motor binding model shows that there is a minimal framework that captures the average system response to increasing turnover.

The feedback relationship from the limited pool of motors guides the core trend in average values. These trends can then be easily picked up by the diffusion model due to the strong linear relationship between density and turnover rate allowing the correlations of the joint distribution to reflect the slope of the trend in average curvature vs turnover. The range of fluctuations in the training data then determines the interpolation distance the model is able to extend these trends across. Much of the predictive power of this diffusion model comes from the fact that the conditioning variable for the model is closely related to the average of a fluctuating data channel.

## V. FUTURE WORK

Expanding the diffusion model to predict sequences of images could allow us to more directly predict cortical flow. Applying this procedure to experimental actomyosin data would be the primary application to fully leverage the predictive power of these models and allow for explorations of parameter space with relatively fewer experimental points. This procedure could also be applied to characterize other systems. Given that long tailed distributions are common in biological data, developing a diffusion model with a long tailed prior distribution, as is discussed in [27, 28], could aid in fully capturing these distributions. Another possible method could be to use active or time correlated noise in the diffusion, which also serves to push values more toward the extremes of the distribution.

Additionally, an investigation of how well the model can infer trends across other control parameters could give more insight into the underlying system dynamics. Considering a parameter without the linear relationship to the data could disrupt the ability of the edges of the distributions to predict new parameter values. However, more spatially organized data could introduce new features for the model to learn relationships from.

## AUTHOR CONTRIBUTIONS

S. V. supervised the research. A. K. B. developed the generative diffusion codebase. Y. Q. developed the Cytosim simulation pipeline. E. R. performed the research and wrote the manuscript.

## ACKNOWLEDGMENTS

We thank Deb Banerjee for useful discussions. S.V. and E.R. were supported by the National Institute of General Medical Sciences of the NIH under Award No. R35GM147400. The authors also acknowledge support from the National Science Foundation through the Center for Living Systems at the University of Chicago.

