## Supplemental Information for "Diffusion models learn underlying trends in actomyosin networks and predict behavior at unseen filament turnover"

### S1. CYTOSIM SIMULATION

The actomyosin network was simulated using the Cytosim simulation package. The simulation is implemented as described in [1] section S1A, with an increased number of motors (4800 initial motors instead of 1600). Actin filaments are nucleated in a 20 x 32 $\mu$m box at a fixed rate and grow at the specified turnover rate for time $t_{grow} = 8.5$s before disassembling. Motors are added to the simulation box at a low background rate of 30 $s^{-1}$. Then, to create the motor gradient driving the cortical flow, they are added at a high rate to the center 10 $\mu$m of the box at an additional rate of 180 $s^{-1}$. Motors are then removed uniformly through the box. Crosslinkers are recycled uniformly across the simulation box. The simulations are run for 100 seconds, recording the system state every second. We do not consider the first 20 s of the simulation run to allow it to reach steady state.

To bin the simulation data into heatmap images, the simulation box is split into bins (1 $\mu$m for the main results). The simulation models actin filaments as a sequence of 0.2 $\mu$m segments. The density is calculated as the number density of segments within the bin, while the curvature is calculated as the inverse radius of the circle through the centers of a given segment and its two neighboring segments.

| | **Actin** | **Myosin** | **Xlinker** |
|---|---|---|---|
| nucleate | $121\,s^{-1}$ | — | — |
| add | 212–740 $s^{-1}$ | 30–210 $s^{-1}$ | $50\,s^{-1}$ |
| remove | 212–740 $s^{-1}$ | $210\,s^{-1}$ | $50\,s^{-1}$ |
| bind | — | $5\,s^{-1}$ | $60\,s^{-1}$ |
| diffusion | — | 10 | 10 |
| unbind | — | $0.15\,s^{-1}$ | $0.05\,s^{-1}$ |
| initial number | steady state | 4800 | 44500 |

TABLE S1. Cytosim parameter values

### S2. DIFFUSION MODEL

The generative diffusion model is an SGM that learn a score function for the diffusion process on heatmap images. The equations for forward and reverse processes are shown in main text Fig 2 respectively given by

$$\begin{aligned} d\mathbf{x}_t &= -\mathbf{x}_t dt + \sqrt{2} d\mathbf{B}_t \\ d\mathbf{x}_\tau &= [\mathbf{x}_\tau + 2\mathbf{S}(\mathbf{x}_\tau, c)]d\tau + \sqrt{2} d\mathbf{B}_\tau \end{aligned} \tag{S1}$$

where $\mathbf{S}(\mathbf{x}_\tau, c)$ is the class conditional score function and $c$ is the turnover rate. The reverse process is solved during sampling using the Euler-Maruyama sampling scheme.

#### A. Architecture

The model uses a NCSN++ style U-Net. We use a base width of 32, with channel multipliers (1,2,4). This results in feature dims of (32, 64, 128) across 3 resolution levels created by 3 residual blocks. Timesteps are embedded into 256 dimensions with a 2 layer MLP, and injected into residual blocks. There are 3 encoder blocks with 3x3 stride-2-convolution, followed by a transformer bottleneck, and 3 decoder blocks with a nearest neighbor interpolation followed by a 3x3 conv. This structure means the bottleneck dimension is image size/4, which for the standard 16x20 image resolution primarily considered in this work results in a bottleneck of 4x5 resolution. Residual blocks use BatchNorm and SiLu activation.

To add conditioning to the model, we considered a few different methods to embed the class label (turnover rate). These included a randomly initialized MLP, mapping turnover rate to a circle, and simply using the class index. Once the label is embedded, it is added as additional channels to the image data, and input into the U-Net without any additional noise. The method of embedding did not result in any significant variation in generated image quality from the model. This is likely because the turnover rate already has an established numerical ordering, so we do not need to learn any semantic relationships between different classes. For all the models discussed in this work we use the circle embedding of $(\sin(c), \cos(c))$, where $c$ is the range of turnover rates normalized to $[0, \pi/2]$, with that range selected to confine the embedding to positive values. We chose this to allow for a null token at (0,0) that is equidistant from all the turnover rates. This feature of the embedding is useful for classifier free guidance (CFG) methods . The general idea of these methods is to combine scores from a conditional and an unconditional model to use both global and class specific features to more fully sample the space [2, 3]. These methods were detrimental for our applications, likely because the predictive power of our model comes from the relationship of the data fluctuations to the control parameter, and the lack of class independent structures in the data make a purely conditional model preferable.

The model is trained on heatmap data from a varying number of turnover rates. To compare across models with varying numbers of seen turnover rates, we use 54,000 training frames evenly split between the rates.

### S3. AUTOENCODER EVALUATION

To identify key features of the simulated heatmap data and develop a metric to compare the high dimensional distributions of heatmap data, we use a convolutional autoencoder to reduce the 16x20x2 heatmap images to a 40 dimensional latent vector. We use 3 convolutional downsampling layers for the encoder, and similarly 3 upsampling layers in the decoder, all with SiLU activations.

### S4. MODEL VARIABILITY

There is some variability in how fully the model learns the underlying distributions, especially for lower number of seen turnover rates. This is illustrated in Fig S1. The model does not converge to a single state even after extended training. Instead after some amount of initial training, the results begin fluctuating around some local minima. Different initializations of training show similar behaviors, but can find different local minima in the landscape. The effect is more pronounced with fewer seen turnover rates, suggesting a loss of detail in the distribution the model is able to learn with larger inference gaps in turnover. This variability in results can be counteracted by training ensembles of different models then selecting the best predictions of the seen turnover rates. This can be measured through the mean squared difference in average curvature and density or through an FID-like metric comparing the Gaussian fits of the simulated and generated distribution in a reduced latent space.

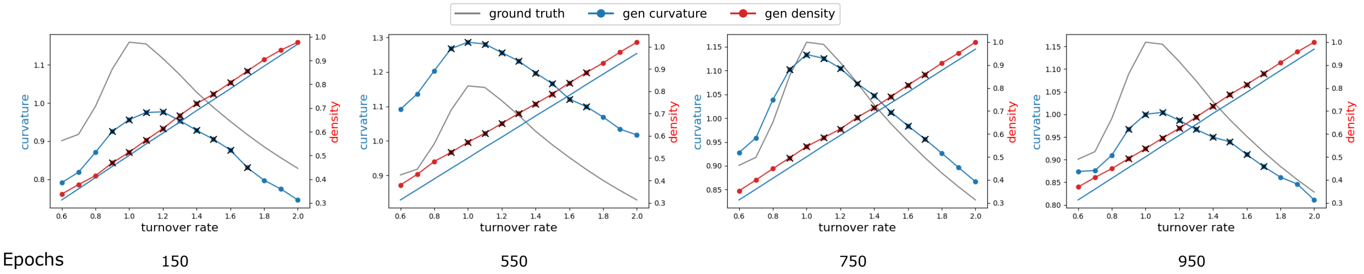


FIG. S1. Illustration of model variability across training epochs

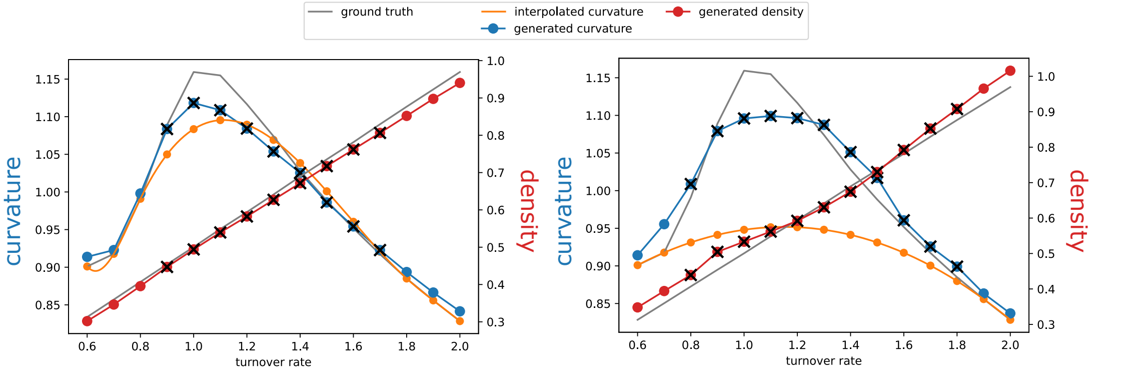


FIG. S2. Comparison of model predictions with naive spline interpolation for 4 and 6 distinct seen turnover rates

### S5. MOTOR BINDING MODEL

#### A. Filament length distribution

A filament grows at rate $k_{\text{act}}$ until it reaches length $L \sim \text{Poisson}(k_{\text{act}}\, t_{\text{grow}})$, then decays at rate $k_{\text{act}}$. Assuming each assembly/disassembly step takes the same expected time, the conditional probability that a filament of maximum length $n$ is in state $l$ is:

$$P(l \mid L = n) = \frac{1}{n}.$$

Marginalizing over $L$ (a filament of length $l$ is only possible if $l \le L$):

$$p(l) = \sum_{n=l}^{\infty} P(L = n)\, \frac{1}{n} = \sum_{n=l}^{\infty} \frac{\lambda^n e^{-\lambda}}{n!}\, \frac{1}{n}$$

*High-$\lambda$ (long filament) limit*

At high $\lambda = k_{\text{act}}\, t_{\text{grow}}$, end effects are negligible, and the distribution becomes approximately uniform:

$$p(l) \approx \frac{1}{\langle L \rangle} = \frac{1}{\lambda} \quad \Longrightarrow \quad l \sim \text{Uniform}(0,\, k_{\text{act}}\, t_{\text{grow}}).$$

$$\mathbb{E}[l] = \frac{\langle L \rangle}{2} = \frac{k_{\text{act}}\, t_{\text{grow}}}{2}$$

$$\text{Var}(l) = \frac{1}{12} (k_{\text{act}}\, t_{\text{grow}})^2 .$$

#### B. Compound Poisson Process for $N_a$

$$\lambda_{\text{act}} = k_{\text{act}}\, t_{\text{grow}} \qquad \lambda_{\text{nuc}} = 2\, k_{\text{nuc}}\, t_{\text{grow}}.$$

The number of filaments $N_f \sim \text{Poisson}(\lambda_{\text{nuc}})$
Total assembled actin, $N_a$

$$N_a = \sum_{i=1}^{N_f} l_i, \quad N_f \sim \text{Poisson}(\lambda_{\text{nuc}}), \quad l_i \sim \text{Uniform}(0, \lambda_{\text{act}}).$$

$$\mathbb{E}(N_a) = \lambda_{\text{nuc}}\, \mathbb{E}(l) = \lambda_{\text{nuc}}\, \frac{\lambda_{\text{act}}}{2} = k_{\text{nuc}}\, k_{\text{act}}\, t_{\text{grow}}^2$$

$$\text{Var}(N_a) = \lambda_{\text{nuc}}\, \mathbb{E}(l^2) = \lambda_{\text{nuc}}\, \frac{\lambda_{\text{act}}^2}{3} = \tfrac{2}{3}\, k_{\text{nuc}}\, k_{\text{act}}^2\, t_{\text{grow}}^3$$

Mean in actin density scales linearly with $k_{\text{act}}$.

#### C. Probability that a motor encounters a filament

Motors are added randomly to the simulation box, and undergo 2d Brownian diffusion. The displacement $x$ at time $t$ is distributed as $P(x,t) \sim \text{Normal}(0, 4Dt)$ The probability that the motor reaches a filament at distance $\xi$ within lifetime $t$ is:

$$P(x > \xi \mid t) = 2\left(1 - \frac{1}{2}\left(1 + \text{erf}\left(\frac{\xi}{\sigma\sqrt{2}}\right)\right)\right) = 1 - \text{erf}\left(\frac{\xi}{\sqrt{8Dt}}\right).$$

The motor lifetime is exponentially distributed with removal rate $k_{\text{rem}} = 210\,\text{s}^{-1}$:

$$P(t) = k_{\text{rem}} e^{-k_{\text{rem}} t}.$$

Marginalizing over the motor lifetime:

$$P(x > \xi) = P_{\text{fil}}(\xi) = \int_0^\infty \left(1 - \text{erf}\left(\frac{\xi}{\sqrt{8Dt}}\right)\right) k_{\text{rem}} e^{-k_{\text{rem}} t}\, dt$$

$$= \exp\left(-\sqrt{\frac{k_{\text{rem}}}{2D}}\,\xi\right)$$

The distance to encounter a filament depends on the network mesh size. Average mesh size scales inversely with actin concentration, approximately as $1/\sqrt{c_{actin}}$

$$\xi = C_1/\sqrt{N_a - C_2}$$

where $C_1$ and $C_2$ are fitting constants to tune the mesh size scaling.

#### D. Motor Binding Rate Equation and Markov Chain

Let $M_b$ = number of bound motors out of total $M$. The mean-field rate equation is:

$$\frac{dM_b}{dt} = -k_{\text{off}}\, M_b + P_{\text{fil}}(\xi)\, k_{\text{on}}(M - M_b).$$

At steady state:

$$M_b^{ss} = \frac{M}{\dfrac{k_{\text{off}}}{P_{\text{fil}}\, k_{\text{on}}} + 1}.$$

To capture fluctuations (rather than just using the average $N_s$), a Markov chain for the full distribution $P(M_b)$ is used:

$$\frac{dP(M_b)}{dt} = -(P_{\text{fil}}\, k_{\text{on}}(M - M_b) + k_{\text{off}} M_b)\, P(M_b) + P_{\text{fil}}\, k_{\text{on}}(M - M_b + 1)\, P(M_b - 1) + k_{\text{off}}\,(M_b + 1)P(M_b + 1).$$


[1] Y. Qiu, E. D. White, E. M. Munro, S. Vaikuntanathan, and A. R. Dinner, Elucidating the Role of Filament Turnover in Cortical Flow using Simulations and Representation Learning (2023), arXiv:2310.10819 [cond-mat].

[2] J. Ho and T. Salimans, enClassifier-Free Diffusion Guidance (2022), arXiv:2207.12598 [cs].

[3] K. L. Pavasovic, J. Verbeek, G. Biroli, and M. Mezard, enClassifier-Free Guidance: From High-Dimensional Analysis to Generalized Guidance Forms (2025), arXiv:2502.07849 [cs].
